# CNSigs: An R Package for the Identification of Copy Number Alteration Signatures

**DOI:** 10.64898/2026.06.21.733646

**Authors:** David Tallman, Shawn Striker, Anjali M. Byappanahalli, Sinclair Stockard, Janet Jenison, Katharine Collier, Elijah Blige, Mark Vater, Daniel G. Stover

## Abstract

Copy number aberrations (CNAs) are gains and losses of large genomic segments present across most cancer types and are a hallmark of cancer genomic alterations. However, the processes underlying CNAs and characteristic patterns of CNAs are poorly understood. Bioinformatic advances have identified underlying single nucleotide variant mutational signatures resulting from distinct mutational processes, yet development of algorithms able to uncover similar signatures for CNAs remains less advanced. Using segmented data files from DNA sequencing, six copy number features are extracted for signature determination: segment size, breakpoints, copy number oscillation, changepoint size, copy number, and breakpoints per chromosome arm, along with ploidy. Mixed model approaches and non-negative matrix factorization are utilized to derive CNA signatures across cancer types. The full methodology was packaged in a publicly available, robust R package, ‘CNSigs’. To verify reproducibility, we derived five signatures from two independent breast cancer datasets (total n>3000), demonstrating high accuracy (average cosine similarity = 0.89). Pan-cancer application of CNSigs in TCGA resulted in derivation of 13 pan-cancer signatures which were significantly associated with disease-specific survival. Benchmarking CNSigs to two other CNA signature approaches within TCGA demonstrated non-overlapping signatures and favorable compute speed for CNSigs. We evaluated n=24 pairs of tumor and circulating tumor DNA (ctDNA) that demonstrated that CNSigs are detectable and reproducible via ctDNA, with significant association of CNSig11 with metastatic triple-negative breast cancer progression-free survival specifically for taxane chemotherapy. CNSigs association with immunophenotype was evaluated in low-grade glioma and CNSig3 was found to be highly prognostic yet complementary to immune features. The CNSigs allows researchers to easily analyze their own samples to derive copy number signatures and evaluate clinical associations. We demonstrate its potential application in ctDNA and association with treatment response. The development of this package allows further investigation of underlying processes that may be responsible for CNA fingerprints.

## INTRODUCTION

Alterations of cancer cells’ DNA is a major hallmark of cancer[1] and DNA alterations are grouped into three main categories: single nucleotide variants (SNVs), small insertions or deletions (indels), or large-scale genomic rearrangement[2]. Somatic copy number aberrations/alterations (CNAs) are defined as either amplification or deletion of larger regions of the genome[3]. CNAs can be pathogenic through gene amplifications leading to overexpression, deletions resulting in lower expression levels or absence of critical genes like tumor suppressors (eg. *TP53*), or impact on regulatory elements (e.g. 3D DNA structure)[3–6]. While specific SCNAs are well-defined, such as the *MYC* amplicon 8q24 altered in ∼10% of all cancers and associated with poor prognosis[7] and marked gene level amplification of the ERBB2 which drives HER2+ breast cancer[8], there are no established tools to efficiently characterize patterns, or “signatures” of SCNAs from tumor sequencing data. Somatic variant-based (SNV) ‘mutational signatures’ can be readily categorized through established computational algorithms[9–11] based on underlying mutational processes, however, few parallel approaches exist for CNAs.

In all cancers, somatic mutations result from multiple mutational processes, such as exposure to mutagens (e.g. UV), defective DNA repair (e.g. homologous recombination deficiency; HRD), and others.[9–11] Signatures of mutational processes are detectable from sequencing data through established computational algorithms.[9–11] Efforts have attempted to develop similar approaches for copy number signatures,[4–6, 12, 13] however, most early approaches focus on one individual copy number feature, such as HRD[13], aneuploidy[4], or tandem duplication[12]. One important study by MacIntyre, et al identified distinct patterns of copy alterations, similar to mutational signatures[14], however, it was only applied to ovarian cancer and was limited in application to tumors with extensive CNAs. In 2022, two tools were released that applied a copy number signature methodology to a wide variety of cancer types in an attempt to find patterns in CNAs that are shared across cancers and relates these patterns back to potential causes of those copy number alterations[15, 16]. However, the implementation of these algorithms remains challenging using publicly-available code, which has led to limited widespread adoption/implementation.

In this study, we present CNSigs[17], a publicly-available, R-based tool that can robustly identify patterns in CNAs across a wide variety of cancer types and settings (**Figure 1**). We demonstrate the application of this analysis pipeline in diverse cancer types, compare the similarities and differences between our method and existing copy number signature algorithms, and apply CNSigs in circulating tumor DNA and in low-grade glioma (LGG).

**Figure 1.**
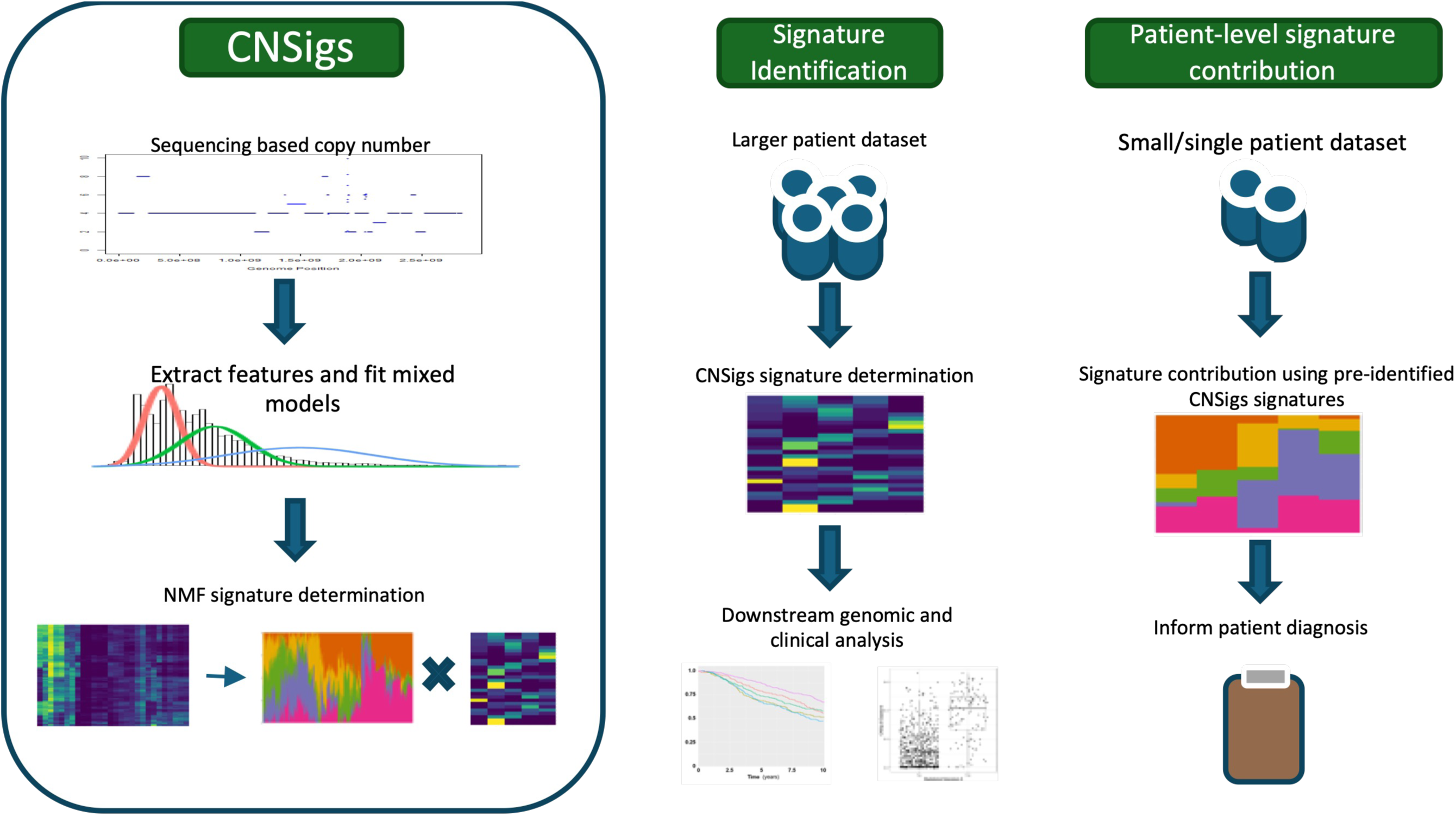
CNSigs Overview. Overview schematic of the CNSigs analysis pipeline and the two main workflows using CNSigs.

## METHODS

### CNSigs Algorithm Development and Pipeline Description

With a goal to develop an easy-to-use, R-based computational package toward the generation of copy number mutational signatures, we developed an analysis pipeline building on the principles used in the original Macyntire et. al study[14] that looked at patterns of copy number alterations in ovarian cancer. In the process, several key changes to the methodology were incorporated into the pipeline to ensure applicability to all cancer types, enhanced reliability across sequencing method, dataset, and the copy number caller algorithm. The overall approach of the pipeline includes: 1) data preprocessing steps; 2) copy number feature extraction; 3) generation of copy number components; 4) dimensionality reduction via non-negative matrix factorization; 5) derivation of new/*de novo* copy number signatures or application of established copy number signatures.

#### Copy Number Segment Smoothing

Copy number segment smoothing is the first step in CNSigs algorithm to ensure artifacts from the copy number callers and sequencing methods are reduced/minimized. This addresses limitations of distinct copy number calling algorithms or sequencing types, for instance lower sequencing depths would often have higher segment sizes. To smooth segments, the copy number profile of sample is assessed for outlier segment artifacts, which are defined as single bin(s) with outlier value in the midst of an otherwise stable segment.

#### Copy Number Feature Extraction

CNSigs algorithm extracts seven different features from the copy number segment data: Breakpoints per 10MB, Oscillations in copy number, Copy number, Changepoints in copy number, Breakpoints per chromosome arm, Segment Size, and Log2 ploidy. Each feature is calculated across the entire genome, except for ploidy, and then is averaged across each chromosome to improve the reliability of the resulting signatures. Averaging reduces the ability for a single feature to dominate the resulting signatures because each feature ends up having the same number of events across the sample; this also improves CNsigs stability when comparing models across different copy number callers. Another benefit of averaging the features across a single chromosome is that it lowers the dynamic range of each of the features and makes the modeling step more consistent and easier to execute, allowing for a single computational pipeline that can be applied to the entire broad spectrum of cancer from genomically quiet cancers to those with a high degree of CNAs.

##### Bp10MB

For this feature, we collapse the entire genome into 10 MB bins. We then look to see whether there is a breakpoint in the sample within the range of each bin. Next, we average the values across each chromosome which results in one value per chromosome, that is the average number of breakpoints seen in a 10MB region for each chromosome.

##### BpChrArm

This feature is simply a count on the number of breakpoint events seen in each chromosome arm. This is the only copy number feature that is not averaged across a single chromosome, and therefore results in two count values per chromosome.

##### OsCN

We define a copy number oscillation as when given a set of three consecutive copy number segments, the first and third segments have the same copy number, that is different from the copy number value of the segment in between them. Therefore, for each chromosome we look to see how many of these oscillation events we see across it, resulting in a single count for each chromosome.

##### Changepoint

The copy number changepoint is the difference in copy number between two adjacent copy number segments. For instance, if the first segment has a copy number value of 2, and the next segment has a value of four, the changepoint between them is 2 (|4-2|). We take the average changepoint value across each chromosome, resulting in a single value per chromosome for each sample.

##### Segment Size

This feature is the length of the copy number segment in bases, calculated by subtracting the start position of the segment from the end position of the segment. The values are then converted to megabase scale (divided by 1E6) so are on a more readable scale. These values are then averaged across a single chromosome, resulting in one value per chromosome for each sample.

##### Copy number

This feature is simply the absolute copy number of each of the segments. The copy number values are averaged across a single chromosome, resulting in one value per chromosome for each sample.

##### Log2Ploidy

This feature is a single value per sample and is appended to the SCM, essentially treating it directly as a copy number component.

#### CNSigs Mixed Modeling Components

After extracting the features, the pipeline then fits a mixture model onto each of the features and uses these mixture models to define the underlying components of the signatures. By default, the package will search for a mixture of between 2 and 10 models of the corresponding distribution types for each feature using the flexmix R package[18–20] (Leisch, 2004). This allows the pipeline to account for cases that have different underlying subgroups for each feature. For the bp10MB, copy number, changepoint, and segment size features, a mixture of normal distributions is fit to the data. For bpchrarm and osCN, a poisson distribution is fit to the data, since both represent count data. Once copy number component models are defined, a sum of posterior probabilities is calculated for each sample per component. The probability that each feature value from each individual sample belongs to each model is determined, then the average of all of the feature values probabilities is determined. This results in the probability that given a chromosome from the patient sample the likelihood that the value for that feature would lie within each of the calculated components.

#### Dimensionality Reduction via Non-Negative Matrix Factorization (NMF)

Using the SCM as the input to the NMF algorithm[21] (with ploidy data appended it to the SCM since it is already a single value per patient), two resulting matrices are generated from the factorization – the signature by component matrix and the patient by signature matrix. The signature by component matrix defines the patterns of copy number features, defining how much each of the modeled copy number components contribute for each extracted signature. The patient by signature matrix defines us how prevalent each pattern of copy number event is in every sample of the dataset, allowing downstream analysis of the copy number signatures.

### Runtime Analysis

We compared the runtime of the analysis pipelines by running each of them 25 times and timing the execution of the full pipelines. We made sure to run each pipeline completely, starting with reading in the data from text files and ending with getting the results. If the pipelines allowed for the use of multiple computational cores for parallel computation, 6 cores were used for all of the pipelines. The analyses were all run on the same computer to ensure a consistent comparison. The specs of the computer were as follows: i7-8700K CPU, 32 GB RAM, NVIDIA GeForce RTX 3090 Ti, Samsung 960 EVO Series - 1TB PCIe NVMe - M.2 SSD.

### Datasets Used in Signature Generation and Interpretation

Patients with breast cancer were identified in The Cancer Genome Atlas[22] (n=725), and METABRIC[23] (n=1424). Samples were included if available ABSOLUTE[24] (for TCGA) or ASCAT[25] (for METABRIC) segmented copy number data and available transcriptome profiling. Established transcriptome-based phenotyping approaches were applied, including breast cancer PAM50 molecular subtyping[26], Single sample Gene Set Enrichment Analysis[27], and TIMER[28]. For comparison of copy number signatures from tumor and ctDNA, segmented copy number data was obtained from Adalsteinsson, et al [29] (**Table 1**). For evaluation of association of ctDNA-based CNSigs with progression-free survival in metastatic triple-negative breast cancer (TNBC), segmented copy number data was obtained from Stover, et al.[30] and Collier, et al. [31].

**Table 1.** Datasets Utilized for CNSigs Tissue versus Circulating Tumor DNA Analyses.

| Dataset name | Sample type | Sequencing depth | # Samples | CN caller |
| --- | --- | --- | --- | --- |
| cfDNA_ULP_WGS_MBC | ctDNA | sWGS (0.1X) | 39 | ichorCNA |
| MBC_WGS_1x | Tissue | 1X WGS | 24 | ichorCNA |
| WES_ABSOLUTE_TM | Tissue | WES (1400X) | 27 | ABSOLUTE |
| WES_ABSOLUTE_ctDNA | ctDNA | WES (1400X) | 39 | ABSOLUTE |
| WES_TITAN_TM | Tissue | WES (1400X) | 27 | TITAN |
| WES_TITAN_ctDNA | ctDNA | WES (1400X) | 39 | TITAN |

ctDNA: Circulating tumor DNA; WGS: Whole genome sequencing; sWGS: Shallow WGS; WES: Whole exome sequencing. All data from Adalsteinsson, et al.[29]

### Statistical Analysis and Data Visualization

Statistical analyses were performed using R version 4.2.0. Data visualizations were performed using ‘ggplot2’[32] and in most cases, with ‘finalfit’ for forest plots[33]. Correlation was calculated using Pearson’s coefficient for normally distributed data and Spearman correlation for uneven distributions. Hierarchical clustering was performed using average linkage. All Kaplan-Meier plots were generated using ‘packHV’ package[34]. Univariate and multivariable Cox proportional hazards models were calculated using the ‘survival’ package in R. All hazard ratios with 95% CI were estimated from Cox regression models.

## RESULTS

### CNSigs Pipeline Usability and Reliability

A core pillar in the development of CNSigs is reliability of results and easy application to diverse datasets. This analysis pipeline has been compiled into a single R package, ‘CNSigs’ that is publicly available on the Comprehensive R Archive Network (CRAN).[35] The package is designed to have as few dependencies as possible to allow installation easily and on many different platforms. CNSigs pipeline allows the user to input the data in several different formats, including a single copy number segment file for all the samples, a path to a folder that has all the individual segment files in it, and an R object if the data has previously been read in. CNSigs allows the user to run the entire pipeline with only a single function call while also allowing fine-tuned analyses on a function-by-function basis. Once results have been generated, additional functions in the package facilitate downstream analyses, including functions for plotting and visualization, comparisons with other CNSigs results, and correlations with other genomic measures.

### Confirmation of Reproducibility: Generation of Copy Number Signatures in Primary Breast Cancer

To confirm the stability of these signatures and to evaluate reproducibility, we applied our ‘signature generation’ approach to two separate, large breast cancer datasets: TCGA (ABSOLUTE-based segments; n=725), and METABRIC (ASCAT-based segments; n=1424). Since both datasets are comprised of similar distribution of breast cancer samples and are both of significant sizes, we would expect that the overall patterns of copy number alterations to be similar. The analysis pipeline was applied to both datasets to generate five *de novo* signatures for both sets of samples, which appeared quite similar in feature contribution (**Figure 2A**). The average cosine similarity between the signatures being 0.91 (range 0.88-0.92; **Supp Fig 1A**). Looking at the per-sample signature exposure across the five *de novo* signatures of each dataset, we can see that the overall distributions of the signatures are similar across all the two datasets (**Figure 2B; Supp Fig 1B**). These data suggests that copy number signatures appear reproducible across dataset and copy number caller.

**Figure 2.**
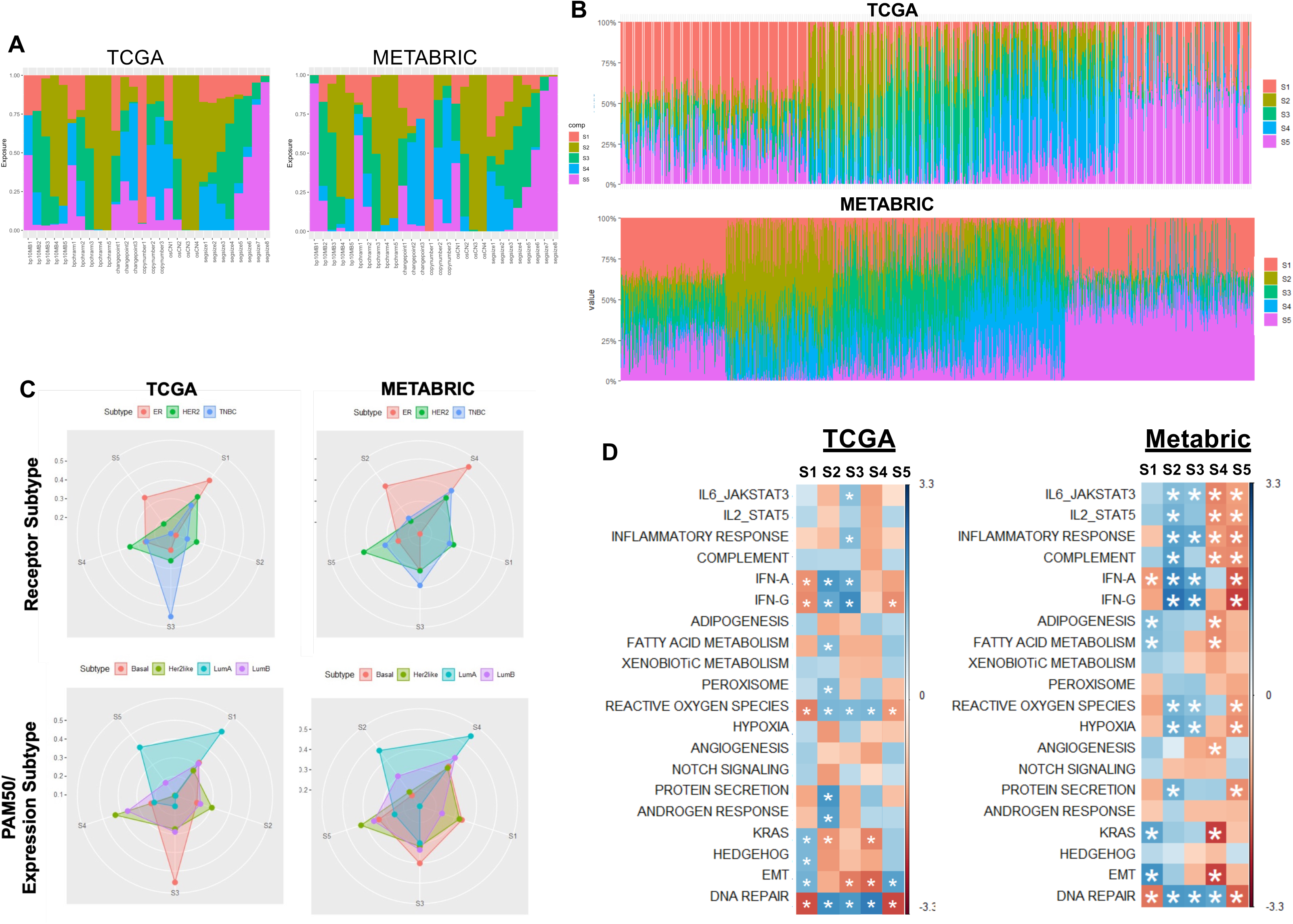
Assessing Reproducibility of CNSigs Signature Generation in Primary Breast Cancer. CNSigs ‘signature generation’ approach was applied to two separate, large breast cancer datasets: The Cancer Genome Atlas/TCGA (n=725), and METABRIC (n=1424). The analysis pipeline generate five *de novo* signatures for both sets of samples with comparable feature contribution (**A**) and per-sample signature exposure **B**). Comparison of exposure of these five *de novo* copy number signatures with known breast cancer receptor subtypes (**C** - top panels) PAM50 expression-based subtype (**C** – bottom panels). (**D**) Correlation plot of exposure to each of the five *de novo* signatures with Gene Set Enrichment Analysis (GSEA) Hallmark Signatures, based on single-sample GSEA for each breast tumor; asterisk indicates FDR p<0.05.

We evaluated the correlation of these five *de novo* copy number signatures with known breast cancer receptor subtypes (e.g. estrogen/hormone receptor positive/HER2-negative; HER2-positive; TNBC) and expression-based subtype (e.g. PAM50) (**Figure 2C**). While the distribution is similar for the two distinct cohorts, each individual copy number signature did not directly overlap with either receptor- or expression-based subtype; rather, each subtype demonstrates exposure of each signature. To assess copy number signatures’ association with expression phenotypes, we determined the correlation of exposure to each of the five *de novo* signatures with Gene Set Enrichment Analysis (GSEA) Hallmark Signatures, based on single-sample GSEA for each breast tumor (**Figure 2D**). While a greater number of the Hallmark Signatures demonstrated significant correlation with the five *de novo* signatures in the METABRIC dataset (likely due to sample number), similar patterns emerged. First, Signature 5 and Signature 1 both were positively associated with interferon, reactive oxygen species, and DNA damage signatures, while Signature 2 was anti-correlated. Signature 3 appeared to be anti-correlated with multiple immune signatures (JAK-STAT, Inflammatory Response, IFNα, IFNγ). Interestingly, Signature 4 showed a strong positive correlation with KRAS and EMT Hallmark Signatures. Overall, this suggests that copy number signatures may be associated with but do not recapitulate established subtypes and gene expression hallmark signatures.

### Comparison of CNSigs to Alternative Copy Number Signature Approaches

While copy number signature analyses are much less frequently applied and less well understood than mutational signature approaches, we did want to compare our CNSigs analysis pipeline to those published by Steele et al[16] and Drews et al.[15] Since all three approaches were derived primarily from the TCGA dataset, we would anticipate at least some similarities in signatures and signature exposure. The current study used the same subset of TCGA used in the Steele et al. paper, which both have an overlap of 6,152 samples with Drews et al., however the underlying copy number features/components vary by pipeline thus it is not feasible to compare signatures directly. However, we evaluated the exposure to the signatures derived by each group (n=13, current study; n=17 Drews, et al.; n=21 Steele, et al.) across the 6,152 shared samples used for algorithm training. Using cosine similarity as the distance metric, it is clear that these three methods share similarities at a high level yet are clearly distinct (**Supp Fig 2A**). Each approach appears to have a subset of signatures that are specific to each approach and do not highly correspond with the others. Conversely, there are several ‘pairs’ of signatures that appear closely related between CNSigs:Drews or CNSigs:Steele yet very few closely correlate between the Drews:Steele methods. There were also a set of signatures from Steele et al. that have a low level of exposure across most of the samples, a pattern which was not replicated in either our analysis or Drews et al. This raises an interesting point about how these two methods may be looking at different phenomena despite both being based on copy number profiles, and that our signatures may be in an intermediate position.

In the development of CNSigs, great effort has gone into usability and runtime as an important consideration. For example, there are several options throughout the analysis pipeline that allow the user to toggle on parallel computing as well as other runtime optimizations. To compare CNSigs runtime with the other published algorithms, a set of 200 random samples from TCGA was used for runtime analysis across 25 repeated runs for each analysis, including CNSigs, Drews, Steele, and sigminer[36], which is an R implementation of the original ovarian cancer pipeline by MacIntyre et al[14]. First, a comparison of runtime for the discovery of a *de novo* set of signatures within the 200 samples, each analysis pipeline was tasked to generate 6 signatures, repeating each run 25 times (**Supp Fig 2B**). The Steele pipeline using CPU averaged ∼125 seconds, while Steele pipeline using GPU and sigminer runtime both averaged ∼100 seconds. CNSigs pipeline showed a consistent runtime of ∼45 seconds, with the Drews (GPU-based) pipeline showing the fastest runtime of ∼30 seconds, yet it was the least consistent with runtimes ranging 18-60 seconds. Next, for identification of signature exposures using a known set of signatures (**Supp Fig 2C**), the sigminer and Steele pipelines had the longest runtime, averaging ∼30 seconds across 25 repeated runs with CNSigs averaging ∼10 seconds and Drews pipeline being the fastest ∼5 seconds. Overall, this shows that CNSigs has competitive runtime, and the extra features to enhance the usability of our pipeline did not adversely affect the runtime.

### CNSigs Pan-Cancer Analysis

Cancers vary widely in the frequency of CNAs by individual cancer type. Using total number of segments, on average we can see that kidney cancers (KICH, KIRC) have relatively few segments while uterine serous carcinoma (USC), sarcoma (SARC), and esophageal cancer (ESCA) have many, though there is variation within individual cancers (**Figure 3A**). To generate a list of copy number signatures across many different cancer types we utilized the TCGA dataset, with 10,674 samples featuring both ASCAT based copy number segment data and ploidy values. All samples from all cancer types were pooled and copy number features extracted using CNsigs. Next, mixed modeling and NMF were implemented to generate a set of copy number components that covered the full range of cancer samples within TCGA, and extracted set of components termed ‘cancerComps’ that was subsequently built into CNSigs package (**Supp Fig 3A-B)**. Using cancerComps, the *de novo* signature analysis pipeline (‘determineNumSigsPipeline’) was applied to each cancer type individually, identifying the patterns of copy number events in each cancer type of TCGA. Each cancer type was run individually to avoid having the resulting signatures skewed towards cancer types that had a larger number of samples within the dataset.

**Figure 3.**
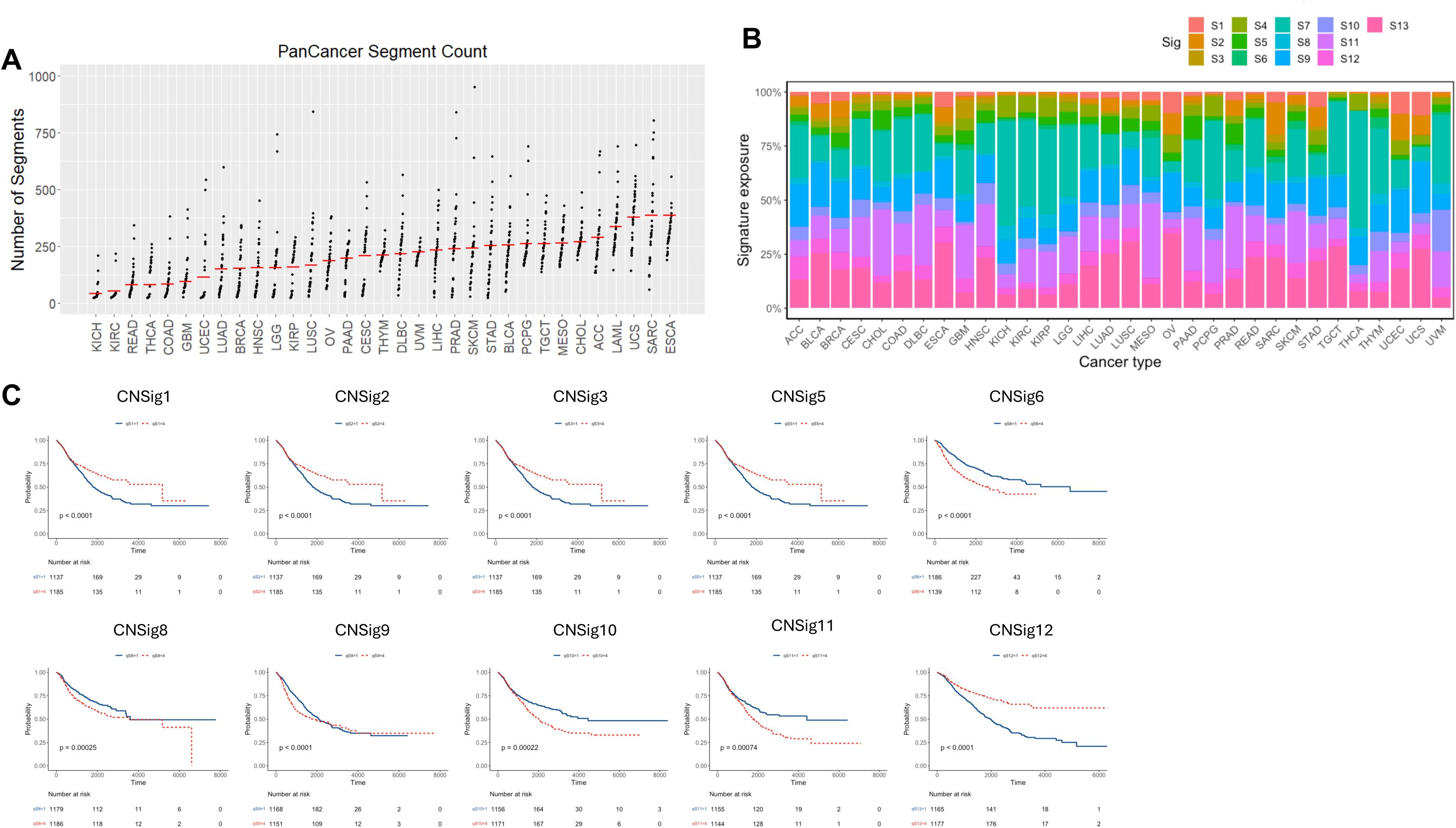
CNSigs Pan-Cancer Analysis. (**A**) Total number of copy number segments, generated by ASCAT, across cancer types of The Cancer Genome Atlas (TCGA). (**B**) CNSigs analysis pipeline applied to the TCGA dataset using the 13 fixed cancer signatures, with average exposure to each of the CNSigs visualized. (**C**) Kaplan-Meier plots evaluating association of CNSigs lowest exposure quartile (solid blue line) to the highest exposure quartile (dashed red line) for each signature across TCGA cancers; nominal log-rank p-value indicated.

To consolidate these 160 cancer type-specific signatures into a smaller number of signatures, cosine similarity was used to compare the contribution of each copy number component from one signature to all other signatures. Using an iterative process to collapse signatures, the two signatures with the highest cosine similarity were merged by averaging the component contribution. This process was repeated until all remaining signatures had a cosine similarity <0.9 with all other signatures (**Supp Fig 3C**). This resulted in a set of 13 unique copy number signatures with distinct feature exposure patterns (**Supp Fig 3D**). The CNSigs analysis pipeline was then reapplied to the TCGA dataset using the 13 fixed cancer signatures (**Figure 3B**). This set of fixed signatures is available in the CNSigs R package[17] as ‘collapsedSigs’ and can be utilized in a similar manner to SNV-based mutational signature analysis.

To evaluate CNSigs performance with different copy number callers, we evaluated the differences in CNSigs output among identical TCGA samples analyzed with ASCAT[25] versus ABSOLUTE[24], two unique copy number callers. Using a dataset of 6861 overlapping samples, the CNSigs pipeline evaluated the exposure of the 13 cancer copy number signatures (**Supp Fig 3B**). The average difference in signature exposure across all of the signatures was 0.02, with the range of signature exposures being 0-1, suggesting robust overall agreement with some samples that showed small differences in signature exposures.

We then evaluated the association of each individual CNSig with disease-specific survival across the entire TCGA cohort (**Figure 3C**), selecting disease-specific survival based on recommendations of reliable outcome data across cancer types[37]. As CNSigs exposure is continuous, to facilitate discrimination in outcomes we compared the lowest exposure quartile to the highest exposure quartile for each signature across available TCGA cancers. Each of the CNSigs is prognostic, though the relative association varies from relatively weak (CNSig9) to early separation and robust discrimination (CNSig12). Similar to assessing whether CNSigs overlapped with existing phenotyping strategies, we evaluated the correlation of CNSigs 13 signatures with SNV-based mutational signatures[38] (**Supp Fig 3E**) but there was little/no correlation suggesting distinct processes as would be anticipated. On the other hand, tandem duplicator phenotype[12], which likely involves double strand breaks, showed a moderately high correlation specifically with CNSigs1 (**Supp Fig 3F)**. Collectively, these data demonstrate that CNSigs can be applied across cancer types and copy number callers and that CNSigs has varying prognostic association when evaluated pan-cancer.

### CNSigs Application in Tumor Tissue and Circulating Tumor DNA

Increasingly, profiling of cancers depends on ‘liquid biopsy, most commonly ctDNA. In advanced cancers there is a larger amount of circulating tumor DNA, termed tumor fraction (or fraction of free DNA in circulation that is tumor-derived) so approaches that leverage genome-level coverage for CNA analyses are feasible from blood. With the rapid and growing potential of liquid biopsy, we evaluated the performance of CNSigs in ctDNA, using existing sequencing data from paired tumor and ctDNA samples, including distinct sequencing approaches (sWGS, WGS, WES), copy number callers (ichorCNA[29], ABSOLUTE[24], TITAN[39]), and samples over time (**Table 1**). Given the potential for low ctDNA content, for this CNSigs application, based on prior work we limited our analyses to samples where ctDNA tumor fraction was ≥10% [40, 41].

First, we evaluated samples with paired tumor biopsy and ctDNA. An overall comparison of CNSigs exposure for tissue (WGS 1X), ctDNA sWGS (0.1X), tissue WES (1420X), and ctDNA WES (1420X) demonstrate generally similar trends of exposure with some variation tissue:ctDNA and sequencing methodology (**Figure 4A-D**, respectively). To furtherevaluate with greater granularity, we visualized paired tumor:ctDNA WES timepoints with copy number segments generated by TITAN for both (**Figure 4E**). While some differences were noted, the agreement was robust with overall agreement (Spearman rho = 0.795). When comparing ctDNA from two distinct time points 2-6 weeks apart, again the agreement was stronger, as might be anticipated (Spearman rho=0.846) (**Figure 4F**). Additional comparisons, including ctDNA sWGS versus WES (**Supp Fig 4A**), tissue WGS versus WES (**Supp Fig 4B**), and paired tumor samples (**Supp Fig 4C**) and agreement was good, yet greater differences were noted when distinct sequencing approaches were used. Overall, CNSigs can be applied to samples across different sequencing depths and coverage, including ctDNA sWGS. The amount of variation captured largely depends on the sequencing depth,; however, the copy number caller can also influence the variation. In general, this supports using a single methodology when applying CNSigs within an individual cohort, for example same sequencing approach/depth and copy number caller.

**Figure 4.**
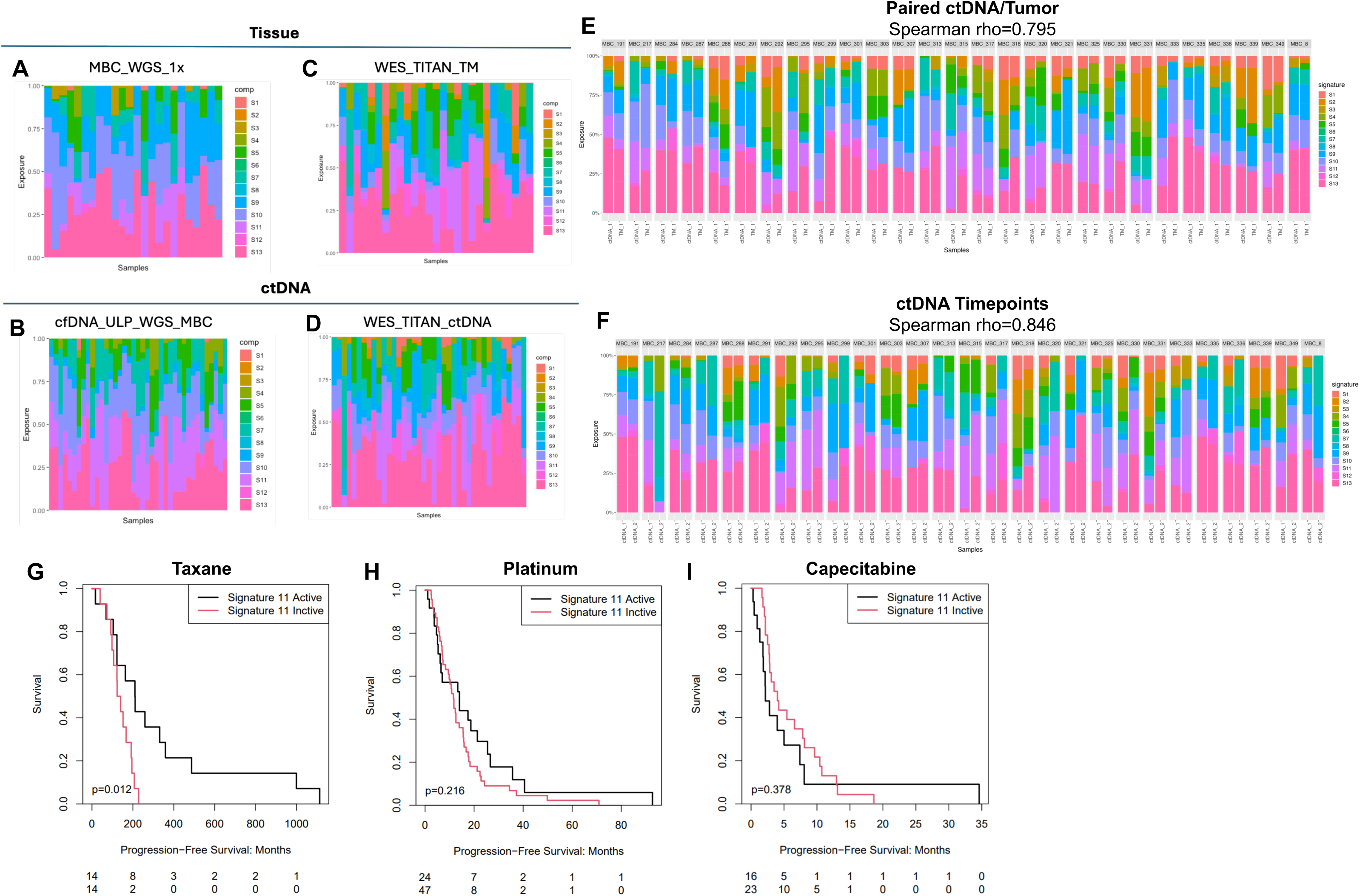
CNSigs Application in Tumor Tissue and Circulating Tumor DNA. Visualization of paired tumor biopsy and circulating tumor DNA (ctDNA; n=24 pairs) per sample signature exposure, including: tumor tissue whole genome sequencing (WGS; median 1X depth) with segments called by ichorCNA (**A**), ctDNA shallow WGS (median 0.1x depth) with segments called by ichorCNA (**B**), tumor tissue whole exome sequencing (WES; median depth 1420X) with segments called by TITAN (**C**), ctDNA WES (median depth 1420X) with segments called by TITAN (**D**). (**E**) Visualization of paired tumor:ctDNA WES with copy number segments generated by TITAN; overall Spearman rho = 0.795. (**F**) ctDNA sWGS (median 0.1x depth) with copy number segments generated by ichorCNA from two distinct time points 2-6 weeks apart; overall Spearman rho=0.846. Evaluation of the association of CNSig11 “active” (exposure ≥0.15; black lines) versus “inactive” (exposure <0.15; red lines) in patients with progression-free survival (PFS) in patients with metastatic triple-negative breast cancer who received taxane (defined as paclitaxel, docetaxel, or nab-paclitaxel; **G**), platinum chemotherapy (defined as carboplatin or cisplatin; **H**), or capecitabine chemotherapy (**I**). Nominal log-rank p-value indicated.

### Circulating Tumor DNA-Based CNSigs as an Actionable Genomic Biomarker

Based on our prior work suggesting one specific amplicon was associated with sensitivity to a specific chemotherapy among metastatic TNBC (mTNBC) patients[31], we hypothesized that specific CNSigs could have predictive potential. TNBC is known to have both extensive CNAs (more than 50% of the genome is altered in most TNBCs) and relatively high tumor fractions in the metastatic setting[30]. To explore the impact of signature exposure on survival, we first determine an optimal cutoff or threshold for a signature being present (rather than just the highest single CNSig exposure). We compared the experimental ctDNA exposures to the paired true tumor exposures using the same threshold value, measuring accuracy, sensitivity, and specificity while sliding the threshold variable between 0 and 1 with steps of 0.05 (**Supp Fig 4D**). To balance true positives and false negatives, a range between 0.15 and 0.25 could be acceptable, although analysis was pursued using the lower end of the range, resulting in a specified threshold value of 0.15 as “active” for survival analyses.

To identify the association of CNSigs with treatment response in mTNBC, we evaluated the association of each of the 13 signatures with progression-free survival (PFS) in mTNBCs with exposure to specific chemotherapy agents: taxane (**Figure 4G**), platinum (**Figure 4H**), or capecitabine chemotherapy (**Figure 4I**). CNSig11 presence (“active”) was associated with significantly longer PFS to taxanes (nominal log-rank p=0.012) but not with platinums or capecitabine. Interestingly, the PFS trend was reversed with capecitabine, with CNSig11 absence (“inactive”) having numerically longer PFS. No associations retained statistical significance after multiple testing correction suggesting that these observations, while intriguing, require further validation.

### CNSigs Association with Immune Features and Survival in Low-Grade Glioma

To evaluate the association of CNSigs with immune features pan-cancer, we interrogated a relatively simple yet consistent measure of anti-tumor immune activity – local cytolytic activity (CYT score or ‘GZPR’), calculated as the geometric mean of the transcript levels of two key cytolytic effectors, granzyme A (GZMA) and perforin (PRF1).[42] We confirmed that the highest quartile GZPR score was associated with improved DSS (relative to lowest quartile) among our TCGA pan-cancer cohort (**Supp Fig 5A**). We then evaluated the distribution of GZPR score across cancer cohorts which demonstrated a wide range in average GZPR score from low (low-grade glioma; LGG) to highest (diffuse large B-cell lymphoma; DLBCL) (**Supp Fig 5B**). When evaluating GZPR score high (above median across entire TCGA cohort) versus low (below median), there were no evident patterns of association with CNSigs (**Supp Fig 5C**). As an alternative approach, we overlaid GZPR score with CNSigs exposure (**Figure 5A**) and noted some intriguing patterns: CNSig7 tended to be over-represented among tumor types with low GZPR score (LGG, PCPG, THCA, GBM), while CNSig11 tended to be over-represented among tumor types with high GZPR score (DLBCL, KIRC, CESC, TGCT).

**Figure 5.**
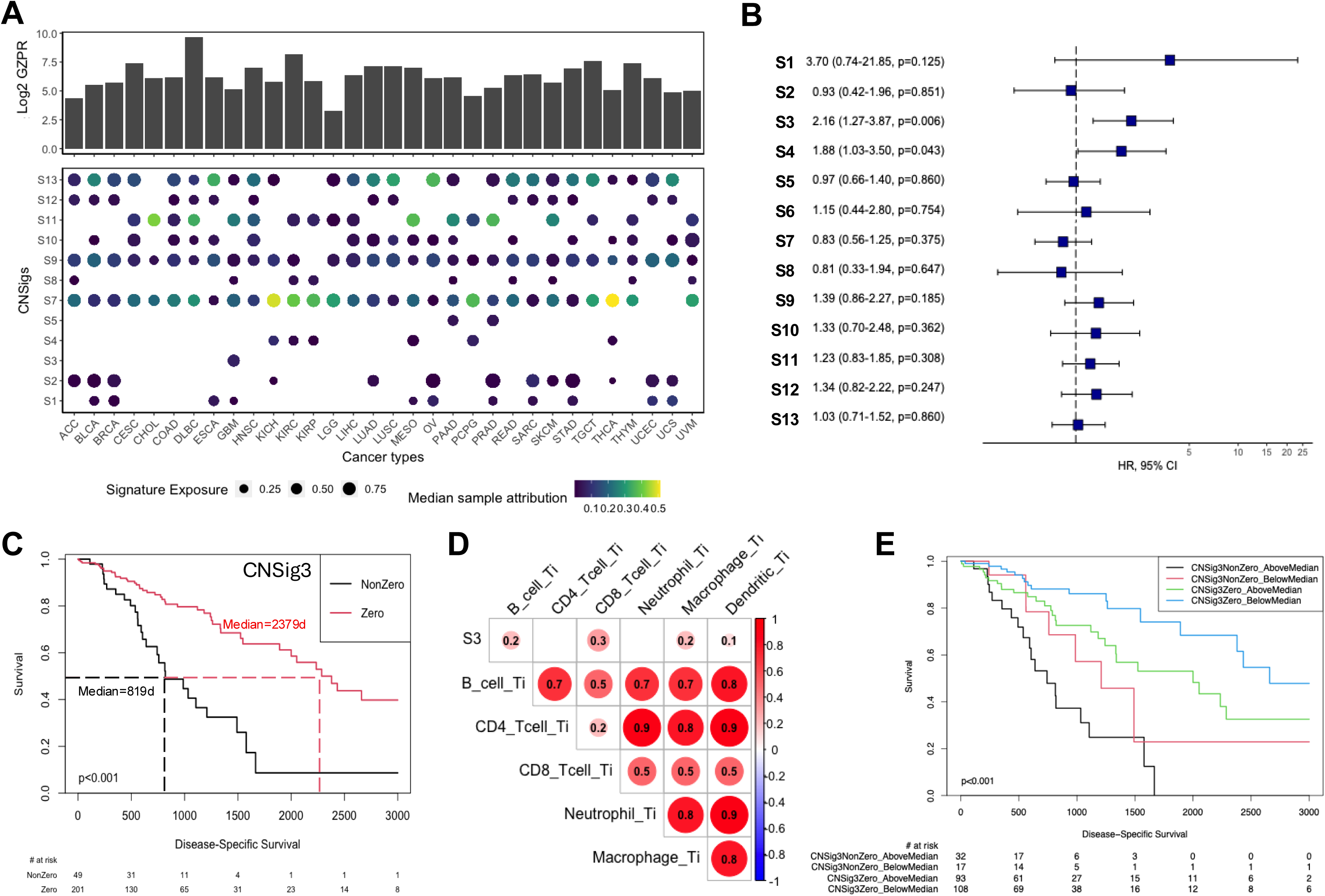
CNSigs Association with Immune Features and Survival in Low-Grade Glioma. **(A)** Local cytolytic activity (CYT score or ‘GZPR’) was calculated as the geometric mean of the transcript levels of two key cytolytic effectors, granzyme A (GZMA) and perforin (PRF1).[42] GZPR score visualized by cancer type within TCGA (top panel) with CNSigs exposure (bottom panel) – size of dot indicative of median sample exposure within cancer type and color of dot indicates median sample attribution. (**B**) Cox’s proportional hazards analysis of CNSig exposure (as a continuous variable) with disease-specific survival (DSS) within low-grade glioma (LGG). Hazard ratio per 0.1 change in exposure, 95% confidence interval, and p-value indicated for each CNSig, visualized as forest plot to right. (**C**) Kaplan-Meier survival assessment of LGG DSS by CNSig3 zero versus non-zero. Median survival for each group and log-rank p-value indicated. (**D**) Correlation plot of Pearson’s correlation of CNSig3 with TIMER inferred immune phenotype. (**E**) Kaplan-Meier survival assessment of LGG DSS by CNSig3 zero versus non-zero and TIMER macrophage above versus below median; log-rank p-value indicated.

We further interrogated LGG, which was characterized by the lowest GZPR score, and found that all 13 CNSigs were represented at varying levels on average among the TCGA LGG samples. When assessing CNSig exposure (as a continuous variable) with DSS by Cox’s proportional hazards, CNSig3 (per 0.1 change in exposure) was significantly associated with worse DSS (hazard ratio/HR 2.16, 95% confidence interval/CI 1.27-3.87, p=0.006) as was CNSig4 (HR 1.88, 95% CI 1.03-3.50, p=0.043) (**Figure 5B**). CNSig3 had zero exposure in most LGG (201/250; 80.4%), thus we evaluated CNSig3 zero versus non-zero. Kaplan-Meier survival assessment demonstrated that CNSigs3 non-zero had significantly worse survival (median DSS 819 days for non-zero; median DSS 2379d for zero; log-rank p<0.001) (**Figure 5C**).

Since LGG showed the lowest average GZPR score among all cancers in TCGA, we then evaluated the association between immune features, CNSigs, and survival. Using RNAseq-based TIMER[28] to interrogate specific immune subsets, within LGG we found that higher inferred macrophage and dendritic cell proportion were associated with worse prognosis, while higher inferred CD4 T-cell proportion was associated with better prognosis (**Supp Fig 5D**). We evaluated CNSigs3 correlation with TIMER immunophenotype, and association was minimal (all Pearson’s r≤0.3; **Figure 5D**), with CD8 T-cells being most strongly correlated. Comprehensively evaluating all CNSigs with GZPR score (**Supp Fig 5E**) or CNSigs with TIMER subsets (**Supp Fig 5F**) similarly demonstrated limited correlation. This suggests that CNSig3 (tumor cell intrinsic) and TIMER subsets (microenvironment) may have complementary prognostic association. To evaluate this, we stratified all LGG samples by TIMER macrophage above/below median integrated with CNSigs3 zero versus non-zero, revealing four groups with distinct DSS, with CNSigs3 zero/TIMER macrophage low having worst prognosis (log-rank p<0.001; **Figure 5E**).

## DISCUSSION

Single nucleotide mutational signatures have had a profound impact of our understanding of carcinogenesis[43–47], DNA repair[48–50], and therapeutic sensitivity[51–54], among other areas. Although somatic CNS remain a hallmark of most cancers[3] approaches to interrogate the patterns or signatures of CNAs are limited. To that end, we present CNSigs as a robust CNA signature algorithm and demonstrate CNSigs utility across cancer types and settings (tumor samples, ctDNA), revealing association with outcome in multiple cancer settings [17].

A critical focus and strength of CNSigs is the broad accessibility and easy implementation. CNSigs usability has been optimized for output from multiple copy number segments algorithms (ABSOLUTE[55], ASCAT[56], and TITAN[39]), and in tumor and circulating tumor DNA. When compared to established CNA signature algorithms from Steele et al[16] and Drews et al.[15], both offer implementation challenges, including multiple different Python analysis packages to execute the full analysis and sparse documentation. Key usability features include: 1) compilation within a single R package; 2) thorough documentation for all functions and overview vignettes that cover the entire pipeline; 3) custom error messages along the entire pipeline to catch common errors; 4) extra functionalities to our analysis package that allow the user to do additional analyses with the copy number data, and allow for several unique and capable data visualizations. The ‘sigminer’[36] package is effectively an implementation of the work from Macintyre, et al[14], which drove the field forward yet is limited in cancer types with few copy numbers. When looking at the runtime of the analysis pipelines, we showed that our pipeline not only had run times that were comparable to all three others while maintaining a high level of usability.

While among the few approaches to computational pipelines to interrogate large DNA alterations in cancer, CNSIgs offers a distinct output for users. Our evaluation of CNSigs relative to Steele et al[16] and Drews et al[15], demonstrated similarities between the signature exposures when applied to the entire TCGA dataset, yet showed that our signatures displayed distinct signatures. Considering other approaches, the ICGC/PCAWG work on structural variations[57] reveals immense complexity yet, requires deep WGS while CNSigs requires only segmented data files from shallow WGS, WES, or even potentially off-target reads from targeted panel sequencing. Other approaches, such as ALPACA (allele-specific phylogenetic analysis of copy number alterations)[58] focus more on clonal evolution than specific patterns of CNAs. Overall, CNSigs targets a middle ground of high usability with sufficient complexity to remain robust.

While SNV mutational signatures have dramatically influenced our understanding of mutational processes in human cancers, they have not yet emerged as a widely used biomarker[59] though promising in clinical management[48]. Similarly, although the initial focus for CNSigs currently is for research application, there is great interest in the potential to leverage CNSigs (and other copy number signatures[15]) to identify biomarkers to guide cancer treatment. We demonstrate robust association with prognosis pan-cancer, although the heterogeneity across cancer types (and even molecular subtypes within each cancer type) limits interpretation. More importantly, we demonstrate both feasibility for CNSigs application in ctDNA – the most common genomic profiling approach for patients with advanced cancer today[60] – and association of presence of CNSig11 with response to taxane (but not platinum or capecitabine) chemotherapy. A broader application of CNSigs across available cancer samples with robust treatment data is underway, which can elucidate potential associations in an unbiased manner. Given the compelling association with microtubule inhibitor taxanes, there are emerging drug classes that target the genomic instability mapped by CNSegs, including separase inhibitors[61], CENP-E inhibitors[62, 63], and cGAS-STING pathway activators (related to release of free DNA out of the nucleus in cancer cells with unstable genomes)[64].

There are limitations to CNSigs and the current study. CNSigs focuses on a subset of large genomic alterations and does not encompass structural variation (for example, work by the ICGC/PCAWG effort[57]). CNSigs does have similarities to existing CNA algorithms[14–16, 36], utilizing NMF-based signature derivation and more computationally innovative neural network-based approaches may offer refined output. While we demonstrate the capacity to apply CNSigs in ctDNA sequencing data, successful application in our experience does require significant levels of ctDNA (e.g. tumor fraction≥10%), which is likely not feasible for many patient samples, limiting broad application. Association with treatment focused on standard chemotherapy agents in the current manuscript, while the breadth of potential treatments is much more expansive.

In conclusion, CNSigs as a robust copy number signature algorithm for cancer researchers compiled into a user-friendly R-package for broad implementation. The copy number signatures generated by CNSigs reflect clinically-relevant association with survival and treatment response.

## SUPPLEMENTARY FIGURE LEGENDS

**Supplementary Figure 1. Supplementary Assessment of CNSigs Signature Generation Reproducibility in Breast Cancer. (A)** CNSigs ‘signature generation’ approach was applied to two separate, large breast cancer datasets: The Cancer Genome Atlas/TCGA (n=725), and METABRIC/MB (n=1424). The analysis pipeline generate five *de novo* signatures for both sets of samples with robust agreement based on cosine similarity. (**B**) Exposure by signature for TCGA and MB.

**Supplementary Figure 2. CNSigs Comparison to Alternative Copy Number Signature Approaches and. (A)** Heatmap comparing CNSigs with the copy number signature approaches from Steele, et al. and Drews, et al. (**B**) Comparison of runtime for the discovery of a *de novo* set of signatures within the 200 samples, each analysis pipeline (CNSigs, Drews, Steele, and ‘sigminer’[36]) tasked to generate 6 signatures, repeating each run 25 times. (**C**) Comparison of runtime for identification of signature exposures using a known set of signatures.

**Supplementary Figure 3. Optimization of Pan-Cancer Signatures and Assessment of CNSigs Overlap with Alternative Phenotyping Strategies.** (**A**) Set of copy number components that covered the full range of cancer samples within TCGA, termed ‘cancerComps’, built into CNSigs package. (**B**) Matrix of the differences in signature exposure across all 13 pan-cancer signatures from TCGA run using either ASCAT or ABSOLUTE copy number callers. (**C**) Cosine similarity plot of 13 unique cancer copy number signatures. (**D**) Feature contribution plots for each of the 13 unique cancer copy number signatures. **(E)** Correlation plot of CNSigs signature exposure with SNV-based mutational signatures.[9] (**F**) Correlation plot of CNSigs signature exposure with tandem duplicator phenotype.

**Supplementary Figure 4. Optimization of CNSigs in Tumor Tissue and Circulating Tumor DNA**. Visualization of including circulating tumor DNA (ctDNA) shallow whole genome sequencing (sWGS) versus whole exome sequencing (WES) (**A**), tissue WGS versus WES (**B**), and paired tumor samples (**C**). Comparison of experimental ctDNA exposures to the paired true tumor exposures using the same threshold value, measuring accuracy, sensitivity, and specificity while sliding the threshold variable between 0 and 1 with steps of 0.05 (**D**).

**Supplementary Figure 5. CNSigs Association with Immune Features.** (**A**) Local cytolytic activity (CYT score or ‘GZPR’) was calculated as the geometric mean of the transcript levels of two key cytolytic effectors, granzyme A (GZMA) and perforin (PRF1).[42] Kaplan-Meier profile of association of disease-free survival among TCGA pan-cancer cohort by highest quartile GZPR score (solid blue line) versus lowest quartile (dashed red line). (**B**) Distribution of GZPR score across cancer cohorts, with plots show estimated value and error bars indicating standard error of the mean. (**C**) CNSigs distribution within each cancer type stratified by GZPR score high (above median across entire TCGA cohort) versus low (below median). (**D**) Cox’s proportional hazards analysis of TIMER[28] inferred immune subsets within low-grade glioma (LGG) and association with disease-specific survival (DSS). Hazard ratio, 95% confidence interval, and p-value indicated for each immune subset, visualized as forest plot to right. (**E-F**) Correlation plot of all CNSigs with GZPR score (E) or CNSigs with TIMER subsets (F) using Pearson’s correlation coefficient.

## Data Availability

All data used in this study are publicly-available and cited within the manuscript.

