## Supplementary figures and images for "CNSigs: An R Package for the Identification of Copy Number Alteration Signatures"

### Supplementary Figures

#### Supp Figure 1

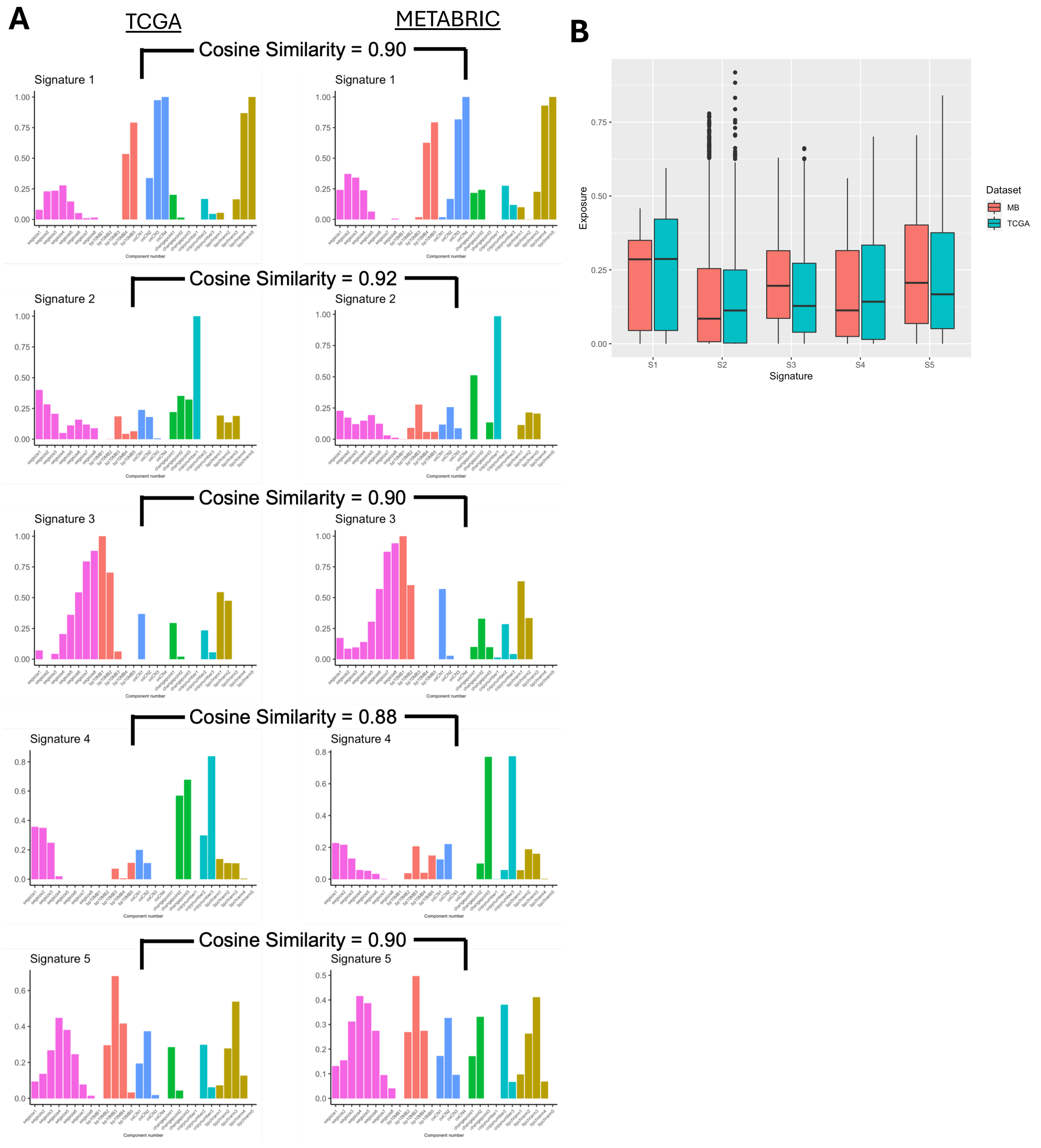

#### Supplementary Figure 2

A

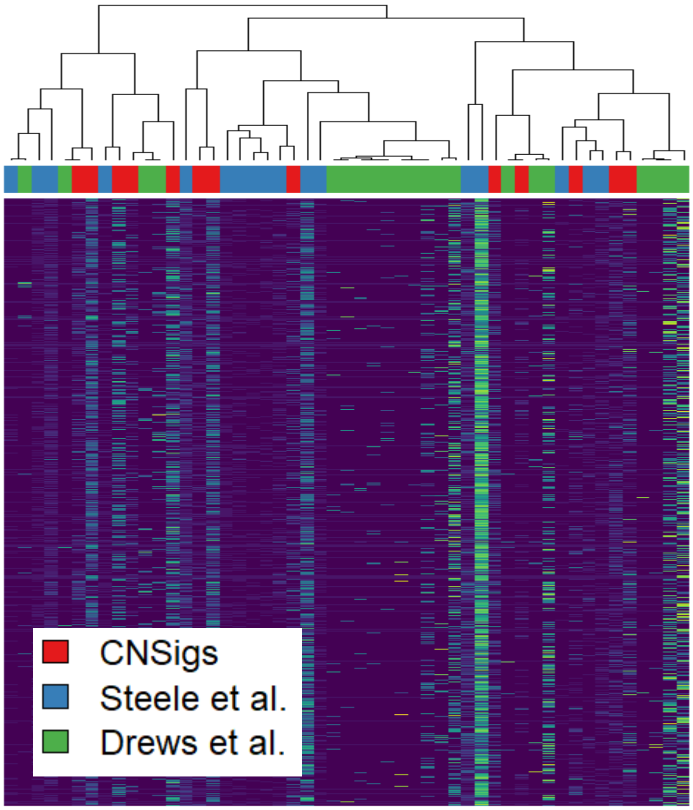


B

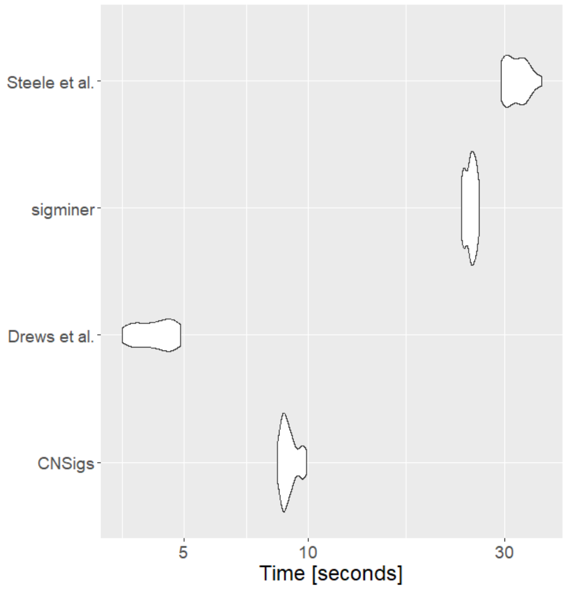


C

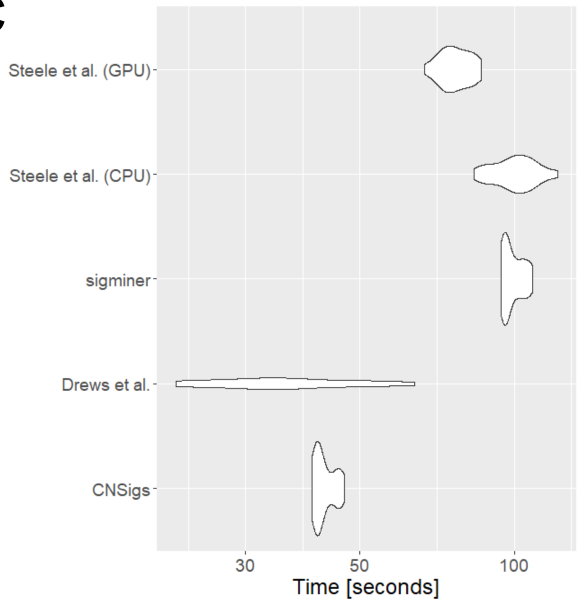

#### Supplementary Figure 3

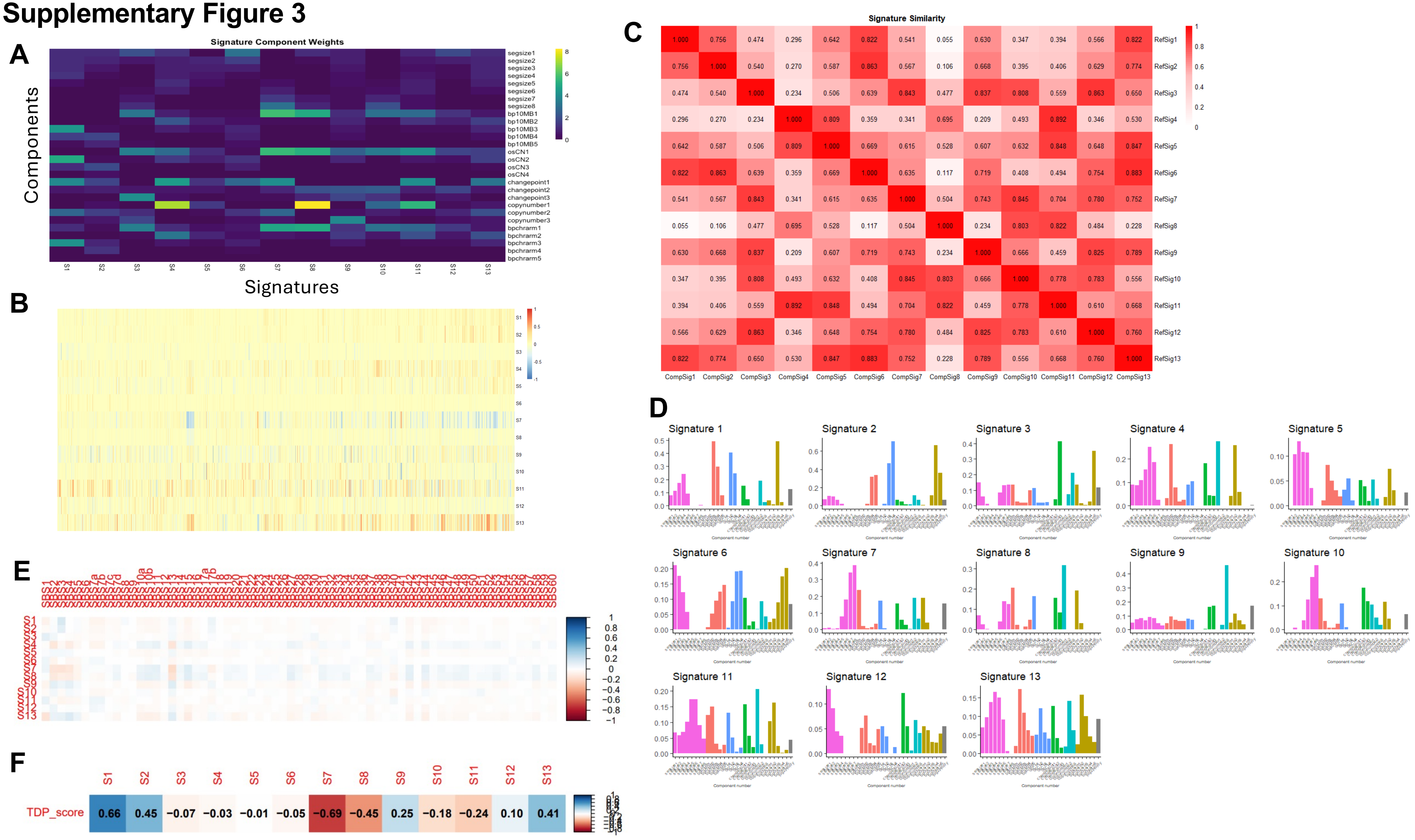

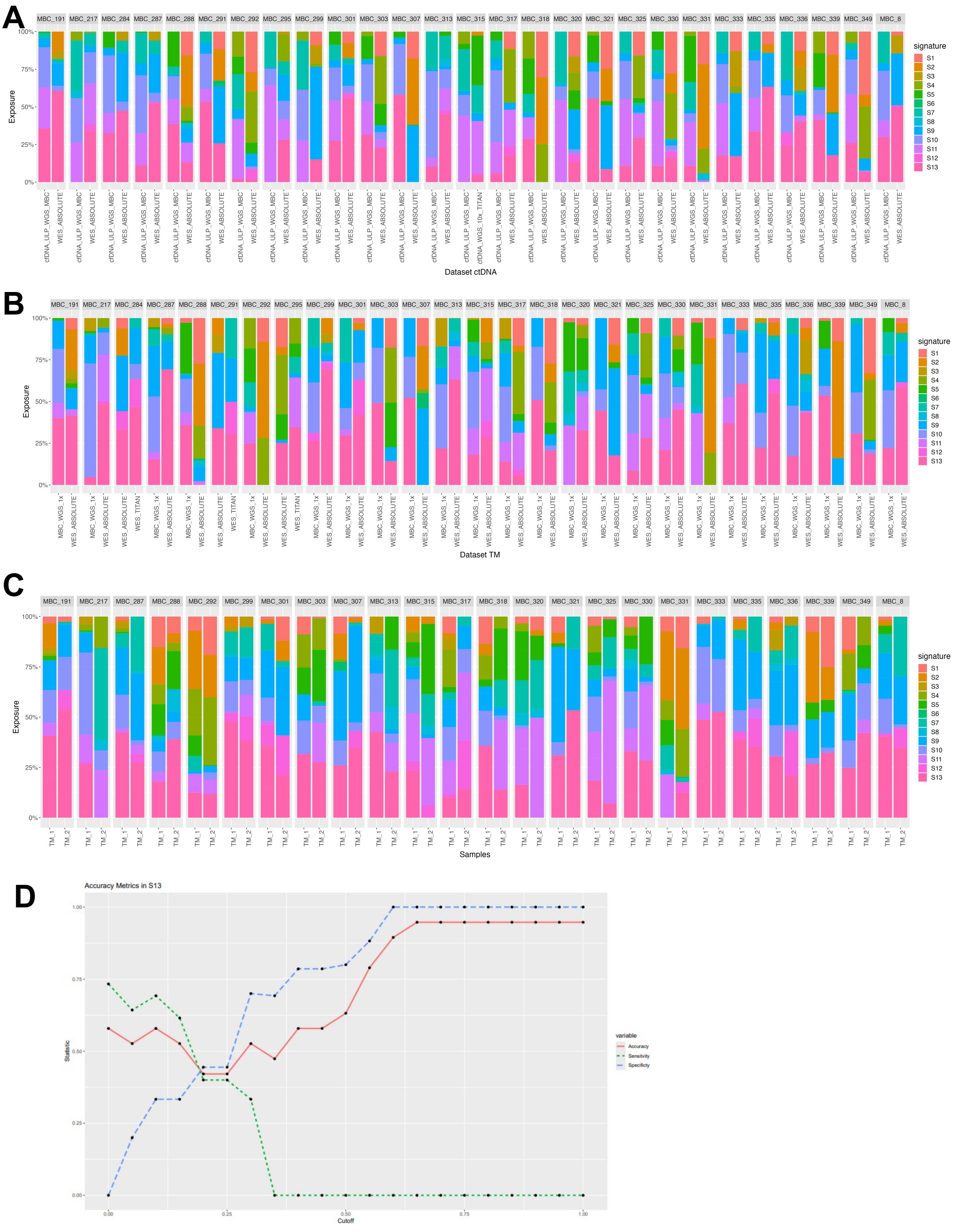

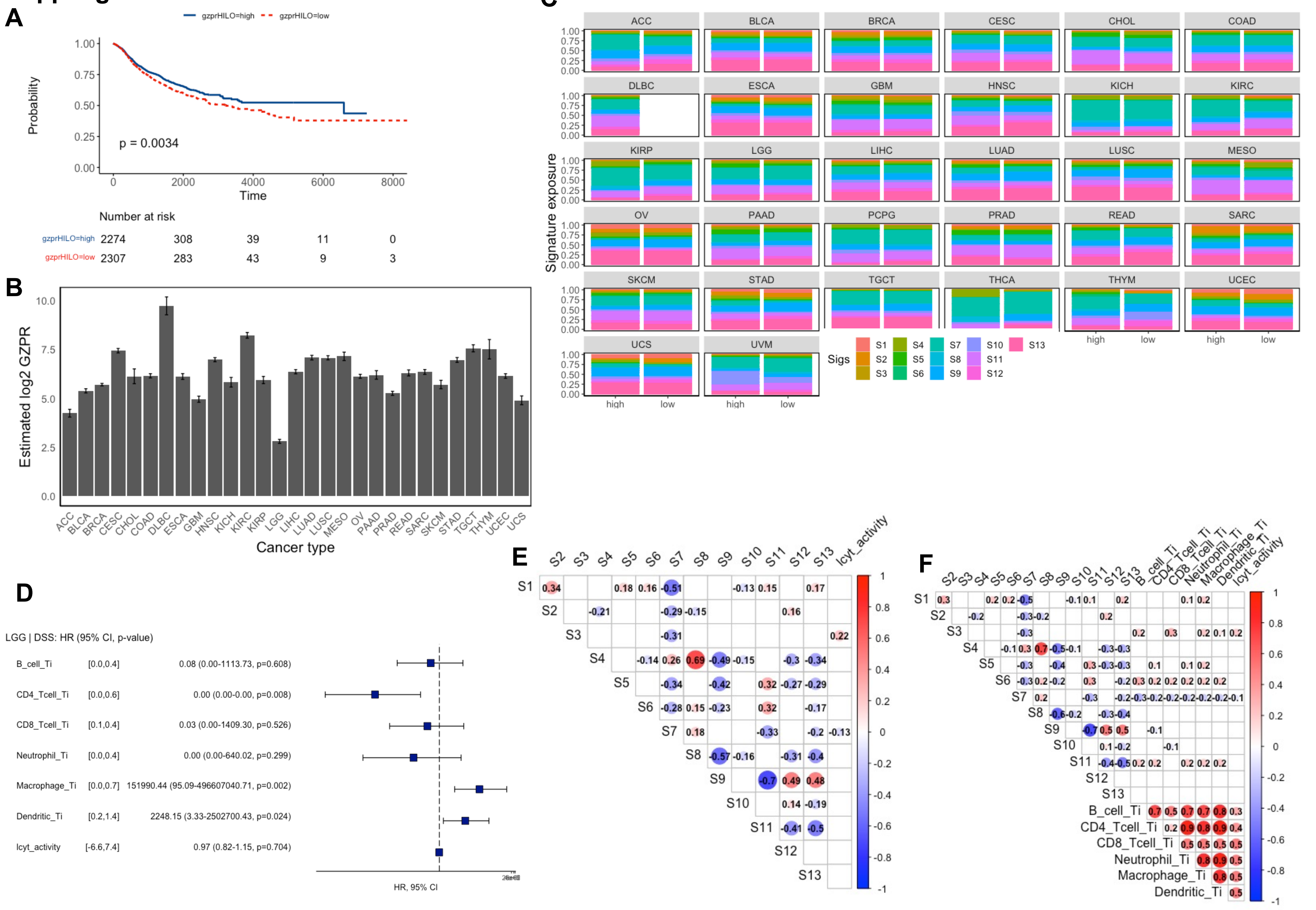
